# The Polymorphisms, Solvent Accessibility and Conservatism of Hepatitis C Virus Nonstructural 5B Protein

**DOI:** 10.1101/2025.02.09.637353

**Authors:** Dejun Lian

## Abstract

The polymorphisms of protein or protein family, that is, the divergences of amino acid and nucleotide sequences have provided much useful information on the divergent evolution of proteins. In this paper, we analyzed the polymorphisms of enzyme NS5B of HCV for which sequence variation among most isolates have been characterized and protein structures of the catalytic domain form of this enzyme are also known. For this protein, we found that solvent accessibility of residues in the protein structure is a strong predictor of whether or not an amino acid will be polymorphic and the residue variability. Apart from polymorphism, we found conservatism at every level among site is universal for this protein. We also found that purifying selection at different levels was strong in the forming of the polymorphisms and conservatism of this protein.

---

The polymorphisms of protein or protein family, that is, the divergences of amino acid and nucleotide sequences have provided much useful information on the divergent evolution of proteins. The neutral theory of molecular evolution posits that the majority of evolution at the molecular level is due to the random fixation of mutations that do not affect fitness. Differences in rates of substitution and levels of heterozygosity among genes, or among different classes of sites within genes, are attributable to differences in the fraction of all mutations that are selectively neutral (mutations that are not selectively neutral are presumed to be deleterious owing to the rarity of beneficial mutations). Selection against these deleterious mutations is known as purifying selection and is acknowledged by most evolutionists as the predominant form of selection at the molecular level (1,2).

RNA viruses, such as hepatitis C virus (HCV), influenza virus, and SARS-CoV-2, are notorious for their ability to evolve rapidly under selection in novel environments. It is known that the high mutation rate of RNA viruses can generate huge genetic diversity to facilitate viral adaptation. RNA viruses offer a unique opportunity for the experimental study of molecular evolution. These viruses exhibit both high replication rates (10^5^day^-1^) and high mutation rates (10^-3^-10^-5^mutation/ (nucleotide /replication)); hence, evolutionary dynamics which would take years to unfold in even relatively simple bacteria occur within days in RNA virus colonies (3,4).

Hepatitis C virus (HCV), a major human pathogen, is a RNA virus which infected approximately 2.5% of the world’s population (130-170 million people) (5). The HCV genome is a positive-stranded RNA molecule of approximately 9400 nucleotides structured in a coding region that contains one large open reading frame and flanked by non-translated regions at the 5’ and 3’ ends. The polyprotein is cleaved into structural (core, envelope 1 and 2) and non-structural proteins (NS2, NS3, NS4A, NS4B, NS5A and NS5B) with one additional small protein at the junction between the structural and non-structural elements (p7protein). The protein NS5B encoded a RNA- dependent RNA polymerase. This enzyme is a key target for developing specific antiviral therapy (6). This RNA polymerase is composed of 591 aa, with a catalytic domain which X-ray structure had been obtained by several groups (7,8).

One characteristic of HCV is its enormous sequence diversity, which represents a significant hurdle to the development of both effective vaccines as well as to novel therapeutic interventions. Phylogenetic analysis of HCV genomes revealed that sequences fall into different clusters. This observation led to a classification of HCV into 8 genotypes and a standardized nomenclature was proposed. The net result of this diversification is the existence of eight major genotypes of HCV that share less than 80% sequence homology with one another, and more than 105 HCV subtypes (9,10). The global distribution of HCV genotypes is regionally specific. Viruses differ from commonly studied organisms such as the geneticist’s mouse or fruit fly, particularly in their speed of sequence change, large population size and the nature of the selection pressures that they encounter. The vast diversity of HCV is mainly due to the error-prone RNA polymerase.

In this paper, we analyzed the polymorphisms of enzyme NS5B of HCV for which sequence variation amongst most isolates have been characterized and protein structures of the catalytic domain form of this enzyme are also known (1). Protein structure acts as a general constraint on the evolution of viral proteins. One widely recognized structural constraint explaining evolutionary variation among sites is the relative solvent accessibility (RSA) of residues in the folded protein. The relative solvent accessibility, which measures the extent to which amino acid side chains are exposed on the surface of the protein or buried within the protein structure, has been shown to predict site-wise evolution in eukaryotes, bacteria and some viral proteins (11). However, to what extent RSA in the protein can be used more generally to explain protein adaptation in other viruses and in the different proteins of any given virus remains an open question.

Apart from polymorphism, we found conservatism at every level among site is universal for this protein.

We also found that purifying selection at different levels was strong in the forming of the polymorphism and conservatism of this protein.

## Materials and Methods

### Sequences and Structures

A total of 12403 sequences of HCV isolates which included all of the eight genotypes were retrieved from the Los Alamos HCV Sequence Database, available at http://www.hcv.lanl.gov/ (12)and Genbank. The nucleotide sequences encoding the protein NS5B were used for comparative analysis. Sequences were additionally filtered that only sequence length are larger than 1000 bases are accounted. The amino acid sequence of 566 aa catalytic domain of NS5B of each isolate were also used for comparative and statistic analysis.

Sites that were variable within species were considered polymorphic, sites that were identical within species were treated as invariant sites. Only amino acids occurring more than once were counted, in order to avoid errors due to a single aberrant sequence.

The structures used in this study were determined through X-ray crystallography. We used PDB code 3I5K structure for analysis because it was more active (13).

### Data analysis

Multiple protein and nucleotide sequences were aligned with BioEdit and edited by hand.

In our analysis, the SAS measure was used to estimate the proportion of each amino acid residue that is accessible to solvent. This was done by taking the ratio of SAS we calculated from the actual protein structure to that of the maximum exposed surface area in the fully extended conformation of the pentapeptide gly-gly-X-gly-gly, where X is the amino acid in question. We then normalized ASA values by the theoretical maximum ASA of each residue (14) to obtain RSA (relative solvent accessibility), We used two methods to estimate solvent accessibility, Solvent accessible surface area (SAS) are calculated using program that implemented in the package MOLMOL (15) and the DSSP program (http://www.cmbi.ru.nl/dssp.html) (16) .The methods gave indistinguishable results.

### Statistical analysis

Data are expressed as means ± SD. Statistical analyses were performed using the Kruskal–Wallis and Mann–Whitney *U* methods. All statistical analyses were performed using SPSS version 13.0 (SPSS Inc., Chicago, IL) with additional analysis performed using Stata/MP14(StataCorp LP). Values of *p* < 0.05 were considered significant.

### Logistic Regression, Confidence Intervals

We used the methods and model of Bustamante CD (1) in understanding how a set of predictor variables affect a dichotomous outcome variable (polymorphic or invariant), the logistic regression is an appropriate statistical model to employ. To assess improvement between nested models that differed in complexity, we used the difference in the log-likelihood of the hypotheses, which is approximately χ2 distributed with degrees of freedom equal to the difference in degrees of freedom of the original models considered.

### The correlation of residue variability with structure

The correlation was tested using Marsh L’s method (17) and Kapp OH’s method (18). I performed several tests to find structural correlates of high evolutionary rate. The correlation of the empirically defined residue differences (where residue differences equals the total number of amino acid states observed at a site minus 1) with structural characteristics was determined to test various models for structural influence on evolution. We used nonparametric analyses (Wilcoxon test, Spearman correlation) because neither amino acid differences nor accessibility are normally distributed. The two-sample rank-sum Wilcoxon test was used to test if groups of sites exhibited significant differences in rates of evolution. Spearman’s rank correlation coefficient, ρ, was calculated to test the correlation between the accessibility of residues and variability.

### The correlation of Entropy at position with structure

To measure the degree of sequence conservation, we calculated sequence entropy for each alignment position within a protein family. Information-theoretical Entropy at position of protein sequences was calculated using BioEdit. Sequences were clustered before calculating to avoid the influence of redundant sequences. We used nonparametric analyses (Wilcoxon test, Spearman correlation). The two-sample rank-sum Wilcoxon test was used to test if groups of sites exhibited significant differences in purifying selection. Spearman’s rank correlation coefficient, ρ, was calculated to test the correlation between the accessibility of residues and entropy.

We used Shannon’s entropy equation which can be formulated as below: H(l) = -Σf(b,l)log(base 2)f(b,l) wheref(b,l)is the frequency of amino acid b(of 20) at the alignment position (19,20) Williamson RM Groups of amino acid physicochemical properties: 20 amino acids are classified into 9 classes by their physicochemical properties as follows:(21–23) 1[V L I M], 2[F W Y], 3[ST], 4[N Q], 5[H K R], 6[D E], 7[A G], 8[P], and 9[C].

H(l) = -Σf(b,l)log(base 2)f(b,l)

Where f(b,l)is the frequency of amino acid grouped .

Using the utilization frequencies of amino acid substitution groups as estimates for their probabilities of occurrence, the information associated with a position is expressed as H (24)

### Ka/Ks

Ka/Ks values were obtained by Datamonkey Adaptive Evolution Server (http://www.datamonkey.org/). The multi-partition fixed effects likelihood (FEL) method implemented in the Hyphy software package on the online server was then used to predict purifying selection (PMID 15703242). HCV sequences were clustered before used.

Ka/Ks values are also obtained by Selection Web Server (http://selection.bioinfo.tau.ac.il). HCV sequences were clustered before used.

### PAM250

PAM250 were calculated by using MegAlign of DNASTAR (DNASTAR, Inc.). HCV sequences were clustered before used.

### McDonald and Kreitman test

McDonald and Kreitman test was done using DnaSP 5.10 program. HCV sequences were clustered before used. We used Canine HCV (Canine hepacivirus AAK-2011 polyprotein gene, complete cds) as outgroup control.

## RESULTS

### Maximum-Likelihood Estimates of Parameters for Logistic Regression of Polymorphism on Solvent Accessibility for Amino Acids Grouped by Protein

Table 1 summarizes the maximum-likelihood estimates of the parameters in the logistic regression model of polymorphism on solvent accessibility for amino acids. The second column lists the proportion of sites that vary within protein. The third and fourth columns give the maximum-likelihood estimates of α and βsas, respectively. The fifth column gives the results of likelihood ratio tests (LRTs) of whether the model with βsas 5 MLE(βsas) fits the data significantly better than a model with βsas= 0, where the test statistic is approximately distributed as χ^2^ with one degree of freedom.

**Table 1.**

| Proteins | N sequences | % Polymorphic | $\alpha$ | $\beta$ | LRT (95% CI) Pr($\chi^2(1)$) |
| --- | --- | --- | --- | --- | --- |
| HCV NS5B (566aa) | 7562 | 95.2 | 0.92 | 3.52 | 32.07 (2.12, 4.92) P<0.001 |
| *E. Coli* & *S. Enterica* Proteins (1955aa) | 138 | 4.5 | -3.87 (-4.26, -3.48) | 3.53 | 33.12 (2.35, 4.71) P<<0.0001 |

NOTE.—CI = confidence interval; LRT = log-likelihood ratio test. a data from ref(1)

**Figure 1.**
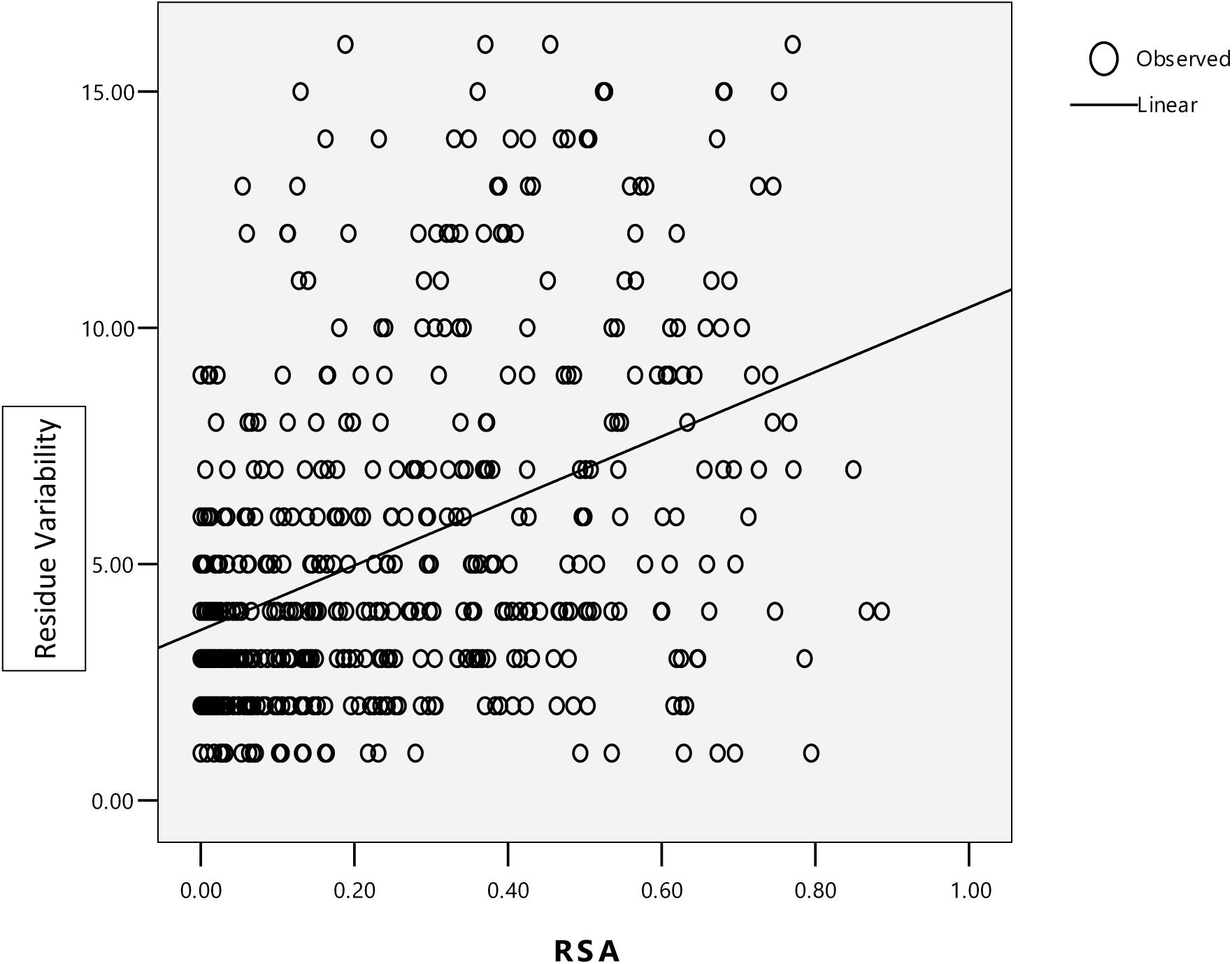
Spearman correlation of residue variability with relative solvent accessibility (RSA) Of HCV NS5B.

**Table 2.** spearman and pearson correlation of RV with SAS.

|  | Spearman 's rho | Pearson |
| --- | --- | --- |
| NS5B (RSA) | 0.420 | 0.424 |
| Rhodopsin (SAS)<br>a | 0.339 | 0.338 |
| Globin (SAS) b | 0.334 | 0.340 |

**Table 3.** Rate Characteristics of Sites in Specific Structural Features of NS5B of HCV.

| Structure<sup>a</sup> | Number of Sites | Differences($\mu_1 - \mu_2$) | Wilcoxon two-sample test |
| --- | --- | --- | --- |
| β-Structure | 61 | 0.61 | |
| α-Helix | 242 | 0.27 | |
| Coil | 213 | -0.35 |  |
| RSA≥5% | 186(35.0%) | | |
|  |  |  | 1.13* |

**\* P <0.01.**
a Designated structural sites compared to remaining protein sites.

### Residue variability

Residue variability, a measure that roughly corresponds to states at residue sites. For HCV NS5B which gave a value of 5.36+_0.14 and increase with RSA The same with Marsh’s result of Rhodopsin. Surface residue 1.13 compare with 1.03 of Rhodopsin More conservative, >=4.7% sequences are monomorphic sites. A great many of the substitutions are conservative substitutions (the same physicochemical class)

### ENTROPY

Average Shannon entropy values were also obtained for individual amino acid positions by using Bioedit Program. The resulting entropy values vary between 0 (total conservation) to 4.332 (complete variation). The mean value of less than 1.0 represent high conservation while less than 2.0 represent intermediate degree of amino acid conversation in the provided sequences (22). For HCV NS5B,which gave a value of 0.422+_0.0216,which is less than 1,indicates high conservation.

Polymorphism and conservatism among site between group has a mean entropy value of0.2859+-0.0167 while the while entropy is 0.4222+-0.0216, showing that there is strong conservative.

**Figure 2.**
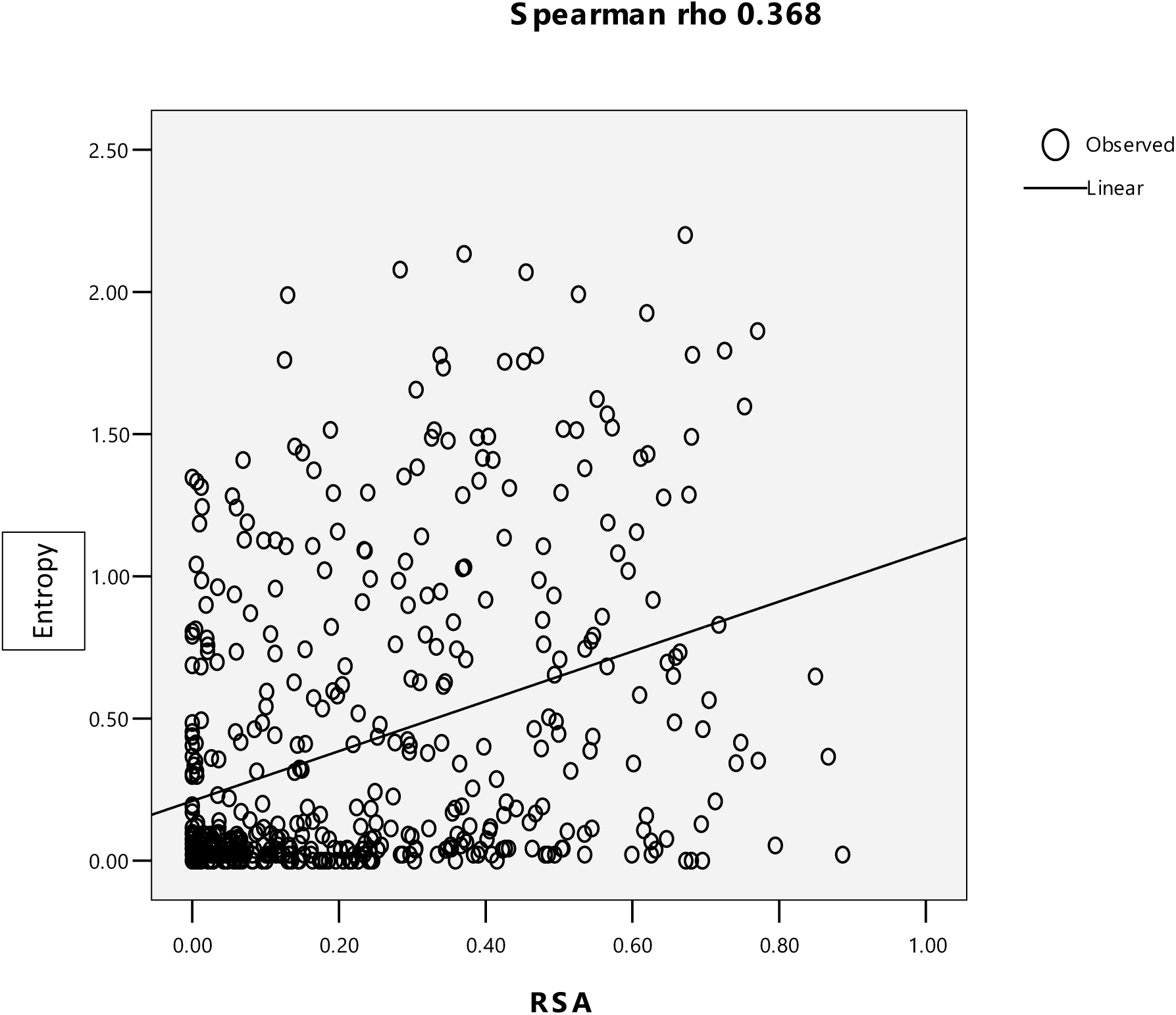
Spearman correlation of Shannon entropy with RSA of HCV NS5B

### Ka/Ks

Classically purifying selection can be inferred by absence. For example, in the Ka/Ks test, we employ the normalized rate of occurrence of substitutions at synonymous sites (Ks) in a protein coding gene as a measure of the background rate of evolution, comparing this to the normalized rate of nonsynonymous changes (Li et al. 1985; Goldman and Yang 1994). A dearth of the latter compared with the former (Ka/Ks < 1) is taken to imply that protein changing mutations happened but were too deleterious to persist (Li et al. 1985; Goldman and Yang 1994).

My result gave HCV NS5B a mean value of Ka/Ks is 0.140+-0.012, which is far less than 1 indicates strong purifying selection over all the protein sequences. Meanwhile, Ka/Ks increases with RSA, showing purifying selection decrease with aa expose to solvent. The relationship between solvent exposure and evolutionary rate (dN/dS) is found to be strong, positive, and linear.in accordance with Franzosa EA and Xia Y’s results.

Table **4** shows the resulting matrix, which represents the first scoring matrix based only on one protein within species.

**Table 4.**
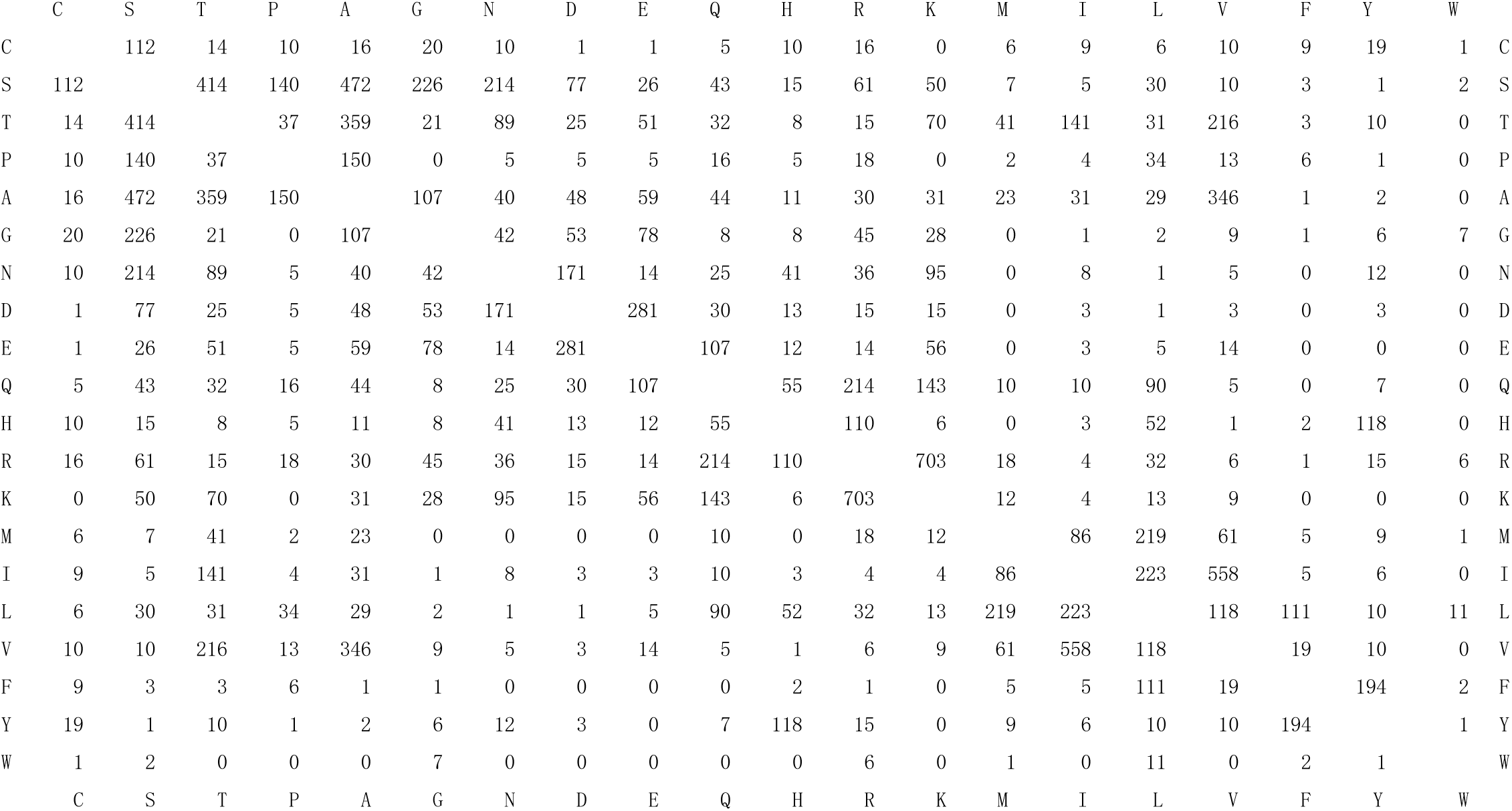
Residue Substitutions of HCV-NS5B Clustal W (PAM250)

### McDonald and Kreitman Table

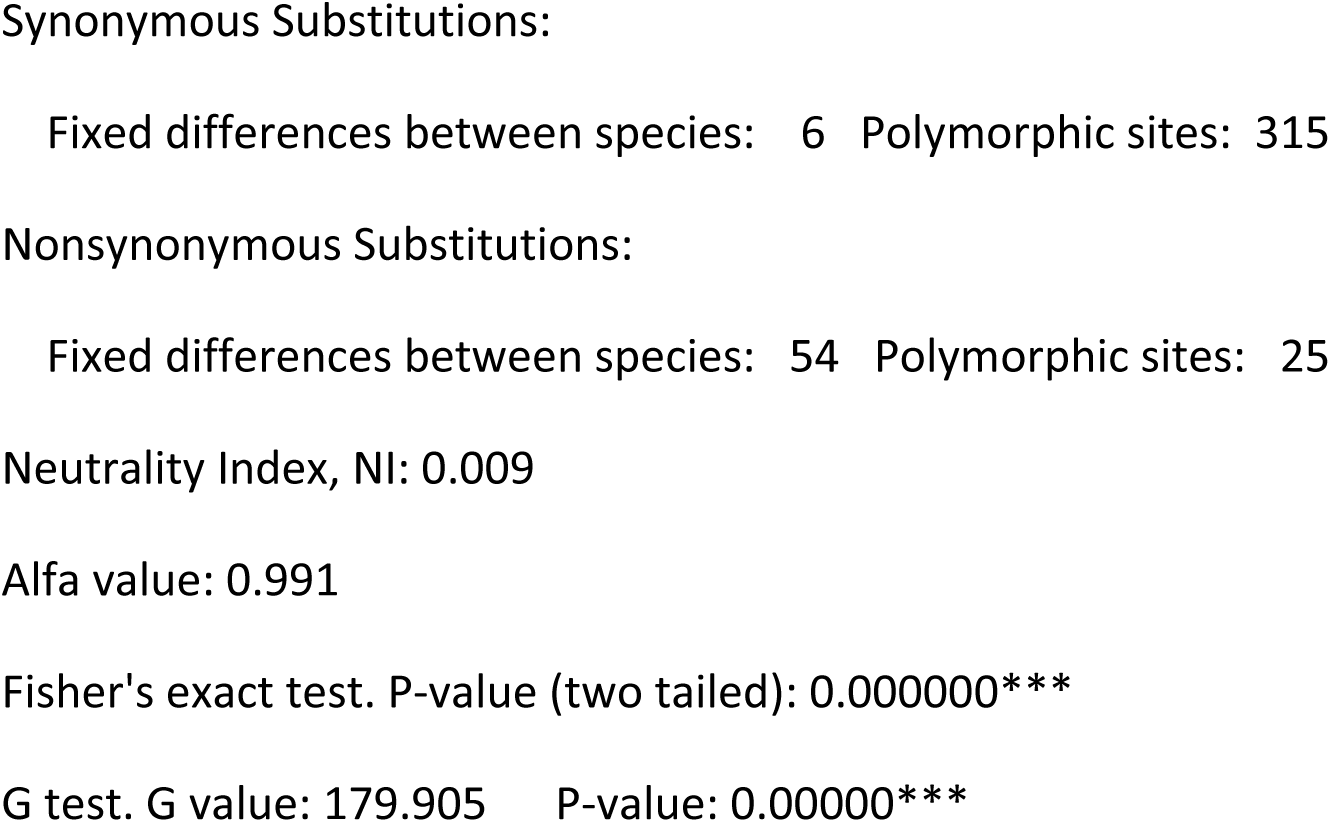
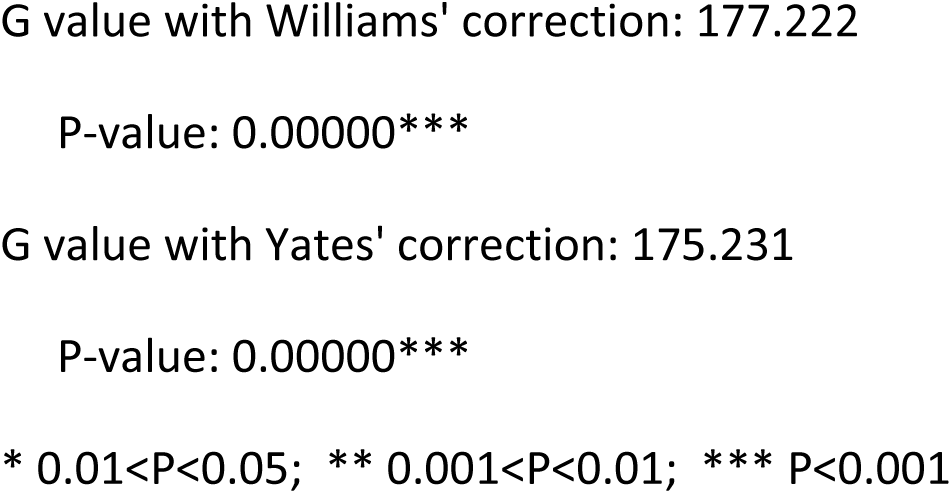

## Discussion

The polymorphisms of amino acid and nucleic acid sequences exist in all protein families. HCV, as a RNA virus which has a polymerase that lacks a proofreading function, shows much more diversity than other species such as *E.coli. S. enteric*.(1) In this paper, I investigated how well different properties of an amino acid—size, physicochemical class, and secondary-structure element—predict the relative likelihood that the site will be polymorphic within HCV NS5B protein. I found a strong positive relation between polymorphism and solvent accessibility, suggesting that amino acid sites that are more solvent-accessible are less likely be constrained in identity. This is the same with Bustamante CD’s result obtained from the study of the polymorphisms of enzymes in *E. coli* and *S. enteric* (1). This finding is in accordance with work done on multiple families of proteins showing that solvent accessibility impact amino acid substitution rates (25-29). This result is also in accordance with the works done on the proteins of yeast (30,31) and many virus proteins (32,33) Unexpectedly, NS5B had a very similar regression coefficient βsas with proteins of *E. coli* and *S. enteric*, suggesting that solvent accessibility may be similarly associated with selective constraints across species differing in myriad details of biological properties and their individual protein structures.

The HCV NS5B protein contains palm, fingers and thumb subdomains. The palm region contains five different motifs A to E, which play a major role in the polymerization ability of HCV polymerase. Motif A contains 212 to 234 amino acids, including the D220-X4-D225 region, which forms the catalytic pocket. D220 and D225 are the residues which are responsible for binding with the magnesium ions. Mutations of D220 to glycine or cysteine completely abolish the NS5B function. Waheed Y’s consensus sequence analysis shows that this region is highly conserved among all the HCV genotypes. Motif B contains G283, T286, T287 and N291 and takes part in sugar selection by NS5B. Their consensus sequence alignment shows that G283, T287 and N291 are highly conserved among all the genotypes while T286 is mutated to proline in genotype 3 and 6. It is reported that the mutation in G283 and T287 completely abolish the HCV NS5B function. Motif C contains the highly conserved GDD motif; the consensus sequence alignment shows that this motif is highly conserved among all the genotypes of HCV. The first aspartate binds the second divalent cation and mutation in this motif is not tolerated, resulting in the abolishment of NS5Bfunction. Motif D contains 326 to 347 amino acids which forms the palm core structure. Consensus sequence analysis shows that R345 is highly conserved among all the HCV genotypes; mutation of arginine to lysine increases the NS5B activity to 152% compared to the wild type NS5B. Motif E contains 360 to 376 hydrophobic amino acids which forms the interaction of palm with thumb.

Consensus sequence analysis shows that this motif is highly conserved among all the HCV genotypes. Consensus sequence analysis shows that the catalytic pocket residues D220, D225, D318, D319, which are responsible for binding with divalent cations, are highly conserved. The other catalytic pocket residues R158, S367, R386, T390 and R394, which interact with NTP triphosphates, are highly conserved among all the HCV NS5B consensus sequences except for the T390 which is mutated to valine in the genotype 5. Global Consensus Sequence of HCV NS5B acid long beta hairpin loop is present in the HCV NS5B protein which protrudes from the enzyme active site. This loop interferes with binding to the double stranded RNA due to steric hindrance. The consensus sequence analysis shows that this loop is highly variable among all the HCV genotypes. It is reported that E18, Y191, C274, Y276 and H502 are important for interaction of template/primer. Consensus sequence analysis shows that these residues are highly conserved among all the genotypes of HCV except for the H502 which is mutated to serine in the genotype 2. Also D225, R48, R158, R386, R394 and S367 are the amino acids which interact with the initiating GTP; previous consensus sequence analysis shows that these residues are highly conserved among all six HCV genotypes (34).

Of the 591amino acids of this protein, only 28 aa are conservative, other 563 aa are polymorphic. Many functional residues like triad GDD are polymorphic. As suggested by Wellner A ‘s result, the laboratory enabled K219S exchange of PGK, which suggested given enough time and variability in selection levels, even utterly conserved and functionally essential residues may change (35).

Although many site-directed mutagenesis studies of several model proteins have demonstrated clearly the ability of protein secondary and 3D structures to accommodate extensive residue variability, the exact limits of the latter have remained a matter of conjecture. Kapp OH’s results (18) for the globin family provide reliable estimates of the maximum residue variability per position, 8-13 overall; furthermore, conservative estimates of residue variability at the interior positions vary tween5 and 7, and these solvent-accessible positions appear to have very little if any restriction in substitution (residue variability 10-15). Our results show that for NS5B of HCV, this is also the case. Reliable estimates of the maximum residue variability per position, 18 overall, Mutation studies have showed that for most of sites of a protein, nearly all of the 20 amino acids can be replaced without significant influence of the function and structure of the protein. While in biological system, this is not the case, there is very strong constraints of residue variability at all sites, which show that many of the replacement may be slightly deleterious and will not be allowed.

Although there is considerable variability of the inner residues (maxium residue variability,11), the inner part of NS5B is very conservative with a mean of residue variability 3.8, (compare with globin family with 5 and 7). Furthermore, part of those solvent-accessible positions appear to have very little restriction in substitution, while others have strict constrains and remain monomorphic. These may all reflect that structural and functional constraints of this protein is very strong both in the inner part and at the surface.

Apart from polymorphism, there are also conservatism of NS5B protein.

Studies also showed that many surface strict conservative residues are involved in enzyme catalysis activity(36,37).There are 28 aa which are completely conservative, they are: 28 N,38 Y,40 T,75 A,102 G,109 R,152 G,192 G,195 Y,200 R,226 S 229 T,265 P,283 G,287 T,298 K,332 D,404 P,410 G,420 W,428 H,489 L492 L,495 P,498 R,503 R,546 D, among them 17 aa are at the molecular surface (This 5% cut-off was devised and optimized by Miller et al. (34), they are:28 N,102 G,109 R,152 G,195 Y,229 T,265 P,287 T,404 P,410 G,428 H,467 H,489 L,495 P,498 R,503 R,546 D, Quite a lot of substitutions the same physicochemical class, which mean structural and functional constraints. show that they are under strong selection pressure. Further studies are needed to test these possibilities.

For decades, rates of protein evolution have been interpreted in terms of the vague concept of“functional importance”. Slowly evolving proteins or sites within proteins were assumed to be more functionally important and thus subject to stronger selection pressure. More recently, biophysical models of protein evolution, which combine evolutionary theory with protein biophysics, have completely revolutionized our view of the forces that shape sequence divergence. Slowly evolving proteins have been found to evolve slowly because of selection against toxic misfolding and misinteractions, linking their rate of evolution primarily to their abundance. Similarly, most slowly evolving sites in proteins are not directly involved in function, but mutating them has large impacts on protein structure and stability (38).

This strict conservation of surface residue may also reflect that there are functional constraints of this protein by the interaction with other proteins. Various studies have showed that protein structure and sequences might be further constrained by selection for interactions among proteins (38). For NS5B, many studies have showed that it has interaction with host proteins (39). There may also be interaction between NS5B with forming RNA replication complex together with other nonstructural proteins and as yet unidentified host components (40). Many of these strictly conservative surface residues may involve in these interactions (39–42).

Another reason may be the need to reduce the stability of incorrectly folded states. Globin and Cyt C. for example, a helix has a hydrophobic patch that ought to pack against another surface, there should not be a competing hydrophobic patch on the wrong side of the helix. This would help to explain why the surface rules are not in effect at all surface sites. and are not very strict even where they are ineffective. Breaking up the continuity of an incorrect hydrophobic patch does not require a rule at every surface site; and a violation might not be too damaging if there are no other violations nearby (43–46).

The amino acid sequences of enzymes like alcohol dehydrogenase and glyceraldehyde-3-phosphate dehydrogenase are strongly conserved across all phyla. Kisters-Woike B et al suggest that the amino acid conservation of such enzymes might be a result of the fact that they function as part of a multi-enzyme complex. The specific interactions between the proteins involved would hinder evolutionary change of their surfaces (47). This may be the case for NS5B protein, NS5B do interacts with other virus proteins and host proteins during infection (39–41).

However, most amino acid replacements with small effects are expected to be deleterious. It has been noted that the low frequencies of most amino acid polymorphisms in natural populations of *E. coli* and *Salmonella enteric* imply that the mutations are slightly deleterious (1), and in the context of the stability-aggregation- degradation model, it is of interest that virtually all of these are physically located in regions of high solvent accessibility on the ‘‘outside’’ of the molecule.

I was interested in a model of evolution in which various types of information such as site amino acid differences, and site structure (access, secondary structure, and environment) contributed to correlations between these observed features were studied to improve predictions of rate for individual sites. Secondary structure type (β-structure, a-helix, and turn did not correlate with amino acid substitution differences). As the result of log regression. residues differences were positively correlated with residue access. Marsh L (18) found no differences in rate among RV with the different types of secondary structure in rhodopsin. However, amino acid differences correlated with residue access (SAS) in TM domains and the whole protein(SAS>70A). Kapp OH (19) found that The residue variability from a subset of 134 monomeric globin sequences was correlated with the mean SAS from nine monomer globin crystal structures. The fitted constants were linear. the correlation coefficients appear to be significant. Sauer RT’s mutation study in biological system (48–51) get the same conclusion with λ repressor using RSA, and roughly liner relationship. My result is the same with them, and do not correlated with Secondary structure elements, and do not correlated with physicochemical property, only hydrophobic residues are slightly less mutable (data not shown), show that the main structural constraint is solvent accessibility (SAS), in accordance with Overington J’s result from work done on multiple families of proteins showing that solvent accessibility has a larger effect on substitution (52). HCV NS5B variability of residues correlated with SAS and has nearly the same correlation efficient with Globin family and Rhodopsin family, showing that there is a common relationship of evolution rate with SAS between different proteins between species and within HCV species.

One measure of such amino acid variation is the Shannon entropy, which is a quantitative measure of sequence dissimilarity that incorporates both the frequencies and the number of variation. Wilcoxin W test showed the entropy surface core difference. The difference between the surface and core reflected that the reduction in purifying selection is so large that sites (near the high end) at the surface of the solvent accessibility range appear to be evolving as fast (median entropy) as those in areas of low solvent accessibility that are under strong selection (median entropy).

In this study I used sequence entropy s(l) as a measure of evolutionary conservation in a protein family. However, multiple sequence alignments used to derive the sequence entropy can be biased (53). Residues can be either invairt (type I), replaced, but physicochemical properties retained (type II). Entropy calculations can be used to identify type I and II, Shen B and Vihinen M use grouped aa entropy to analyze type II conservation (18). Mirnyet Let al. use grouped entropy to study evolution conservation (54). I found that Shannon entropy of NS5B strongly correlated with RSA, show that the outer side of the molecular is less conservative. Futher more, I found that the grouped Shannon entropy reduced dramatically with group, which show conservative within aa physicochemical properties.

Studies from protein families show that amino acid substitutions among physicochemically similar amino acids are more frequent than those between dissimilar ones. I find this kind of purifying selection for protein polymorphism act at conservative amino acid level –Constraints on sequence diversity-purifying selection.

Analyss of available HCV full genome sequences from public databases shows that the degree of sequence variation varies both between different proteins, but also between regions of the same protein. Some regions are highly conserved even across different HCV genotypes (data not shown). Many of these highly conserved regions represent functionally important motifs in the viral protein in which substantial sequence variation is not tolerated. Viral evolution is clearly limited by structural constraints forcing the virus into a state in which it is able to functionally exist. Many mutations that occur during the replication process are deleterious or disadvantageous to the fitness of the virus and are, therefore, negatively selected. This kind of purifying selection is a main selection force for NS5B.

The Ka/Ks value of the whole protein indicate that there is a strong purifying selection in this protein, in accordance with previous studies (55,56). The correlation of Ka/Ks with RSA is used in conjunction with data on synonymous polymorphism to estimate quantitatively the reduction in purifying selection with increasing solvent accessibility. When compared with the distribution of synonymous polymorphism, the increased probability of amino acid polymorphism with solvent accessibility (fig. 3) suggests strong purifying selection in areas of low solvent accessibility and weak purifying selection in areas of high solvent accessibility, irrespective of synonymy class.

**Figure 3.**
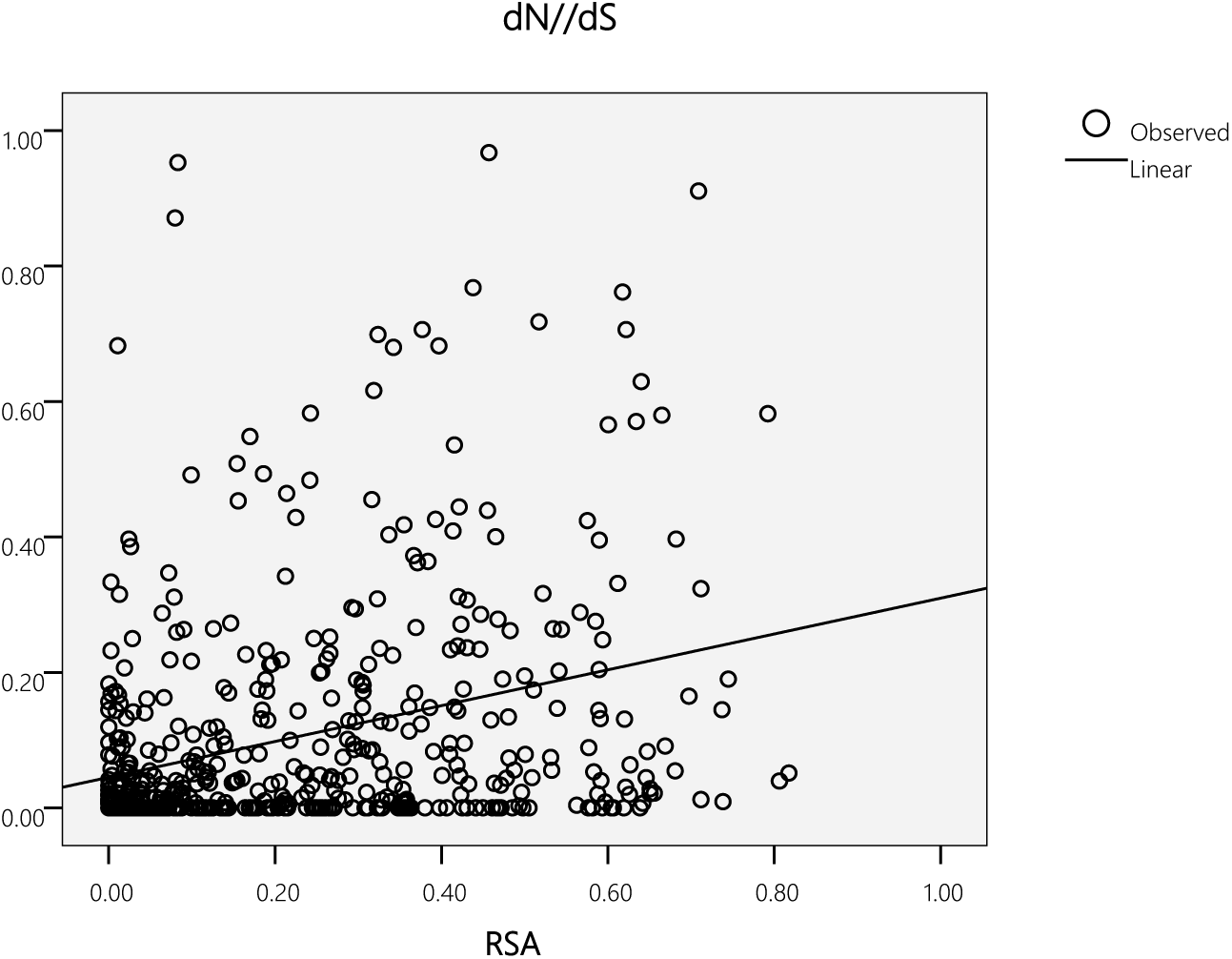
correlation of dN/dS values with RSA

**Figure 4.**
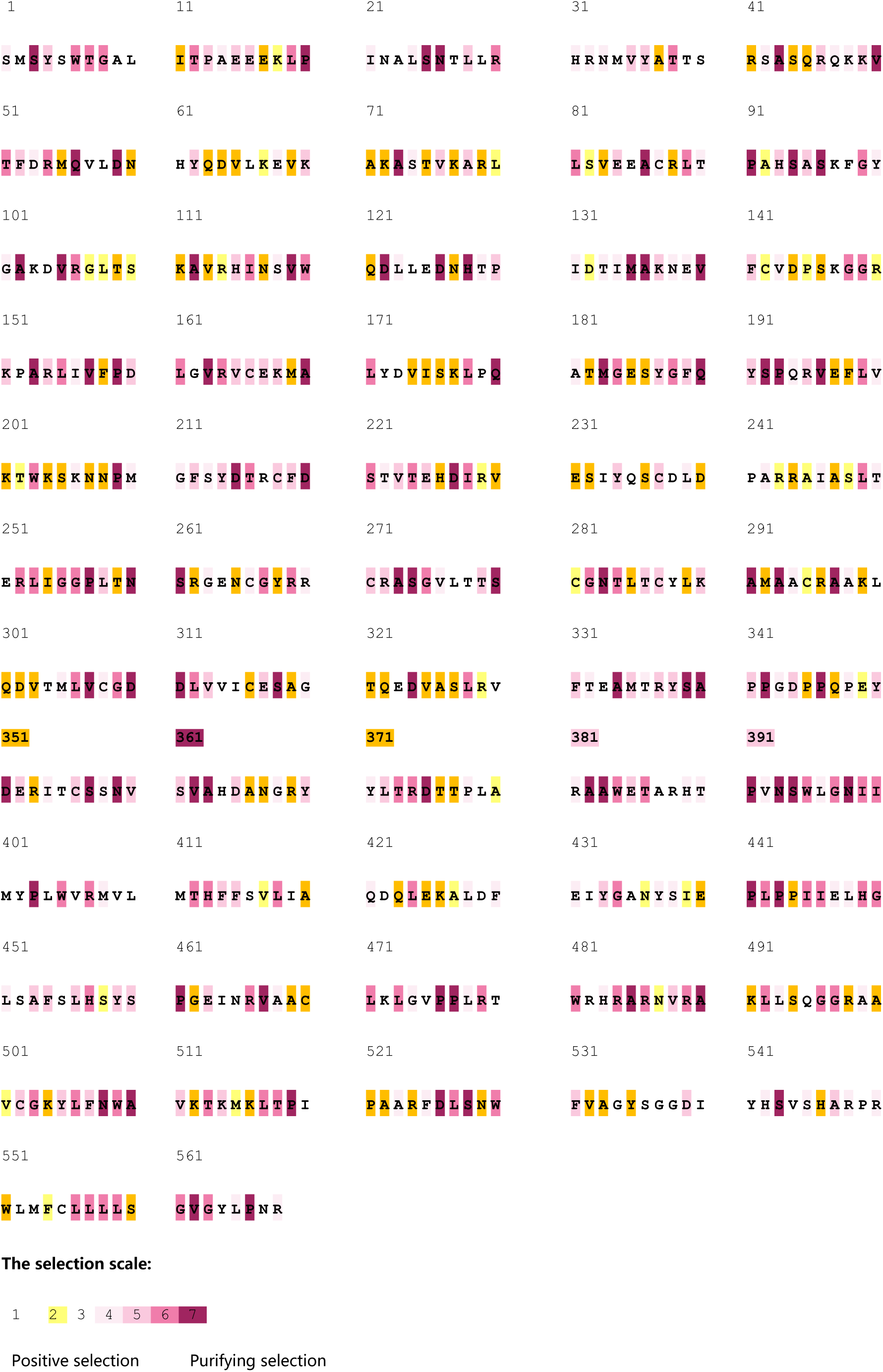
Ka/Ks values of HCV NS5B.

The relatively high Ka/Ks values of this protein reflect the high mutant rate of this protein and also the saturation property of its coding sequence at synonymous sites. For NS5B, Nearly all of the 3ed position of the coding sites are saturated for both twofold and fourfold redundant sites(data not shown), only at 21sites (3.6%)) the synonymous diversity are suppressed due to virus RNA secondary structure proposed by Simmonds P(57).There are also saturation phenomena at synonymous sites in genes encoding protein families between species which may reflected strict constraints at those sites (58).This saturation also means that purifying selection may be stronger than expected form the Ka/Ka value.

The Ka/Ks value of two-fold redundant sites at the surface is smaller than that of proteins of *E. coli* and *S. enteric*(data not shown), this may reflect that purifying selection in HCV may be stronger than that in *E. coli* and *S. enteric*, at least at the surface of NS5B.This may be due to the reason that many of its surface residues are functional important and remain conservative; and there is strong purifying selection by the constraints of interaction of NS5B with other host proteins and other virus proteins (39–41).

In contrast, multiple forces of the immune system or in some cases drugs exact positive selection pressure away from the consensus sequence in the individual. Selection of variants is, therefore, a trade-off between host pressure and functional needs. Purifying selection describes the driving force towards a sequence with optimal replication capacity in the absence of outside pressure on the virus. Reversion of resistance mutations that have been selected in the presence of antiviral drugs back to the consensus sequence have been first described in the influenza model. For HIV and HCV, reversion of drug resistance mutations is well documented and the concept of a salvage therapy was based on this observation, in this concept treatment is re-initiated after interruption and genotypic reversion of drug resistance. This kind of selection show that strong purifying selection for protein polymorphism act at conservative amino acid level (59–61).

McDonald and Kreitman test showing that the Neutrality Index, NI: 0.009(G test. G value: 179.905 P-value: 0.00000), which is far less than 1 which indicate that an “excess” of amino acid evolution between species(63) .Polymorphism data for NS5B genes from both Canine HCV and humans HCV show a significant excess of amino acid polymorphism relative to that expected from the patterns of divergence at silent (synonymous) and replacement (nonsynonymous) sites (62). They show an excess of amino acid polymorphism over that expected from a strictly neutral model. The frequencies of silent and replacement polymorphisms may also shed light on the mildly deleterious model of evolution. While certain “silent” sites are undoubtedly under some functional constraints, which show that purifying selection is strong for HCV NS5B。

The evolutionary divergence of protein amino acid sequences is subject to purifying selection against amino acid substitutions imposed by functional and biophysical constraints. Due to such constraints, the sites (residues) of a protein amino acid sequence differ in their evolutionary rate (the number of amino acid substitutions per unit of evolutionary time). As a result, multiple alignments of evolutionary related (homologous) proteins show clear site-dependent conservation patterns. Typically, only a few sites are directly related to function and their high conservation is due to direct function-specific selection. Mutations at most other sites affect fitness indirectly through their effect on the protein’s folding, stability, structure, or dynamics . (64–66).

The studies on the evolution of protein families with their tertiary structures show that because many differing amino acid sequences are able to attain the same three-dimensional arrangement of the polypeptide main chain, one may conclude that there may be little restraint on amino acid sequences over long periods of time (48). For NS5B of HCV as a within species protein family, the situation is different. Despite its great diversity, the amino acid sequence of this protein is very conservative compare with those between species protein families such as globins and rhodopsins. DuBose RF *et al ‘s w*ork on the polymorphisms of Bacterial Alkaline Phosphatase of Enteric Bacteria showed that most of replacement was conservative and the DNA sequence of silent sites were saturated (67). Sawyer RT and Hartl DL’s work on Drosophila using a Poisson random field model showed a very low value of the replacement mutation rate, relative to the silent mutation rate, may reflect mutations occurring in only a small number of codons at which **a** favorable amino acid change is possible, with all other changes being strongly detrimental (68). So that there is a strong conservatism of amino acids of proteins within species.

This conservation of protein sequences may be a general phenomena of within species protein families which reflects strict structural and functional constraints and other constraints within species. As in the study of human SNP. The mean residue variability 3.6 with a standard Deviation of 3.1, which corr0.14--0.289, they estimate that 50% of mutations are likely to have mild effects, such that they reduce fitness by between one one-thousandth and one-tenth. they also infer that 15% of new mutations are likely to have strongly deleterious effects (68–70). They estimate that 19% and 20% of nonsynonymous mutations are neutral is remarkably consistent (although somewhat larger) than Eyre-Walker A et al.’s estimate of 23% (21–29%) inferred from a smaller dataset using folded frequency data and the hybrid approach of Yampolsky LY et al. expect a slightly larger proportion of mutations to be nearly neutral, they infer that 19% of mutations are effectively neutral and that 14% of mutations are slightly deleterious, such that they segregate in the population at moderate frequencies, but never become fixed. The remainder of the mutations are strongly deleterious such that they contribute little to polymorphism or divergence (71,72). I found that 19% of mutations are neutral or nearly neutral.

Exposure to solvent is one such structural property that has received attention as a correlate of molecular evolution, both at the whole-protein and residue levels. It was observed early in the history of structural biology (73) and has been repeatedly confirmed since, that residues buried in a protein’s core are more likely to remain conserved during evolution than their solvent-exposed counterparts. Dai L et al showed that for NS5A protein of HCV, the distribution of fitness effects of mutations is modulated by both the constraints on the biophysical properties of proteins (i.e., selection pressure for protein stability) and the level of environmental stress (i.e., selection pressure for drug resistance, they found that the fitness effects of deleterious mutations at buried sites (i.e., with lower solvent accessibility) were more pronounced than those at surface-exposed sites(74). This was the case for HCV NS5B protein.

Hartl DL had studied the polymorphism of amino acids based on frequencies alone using Poisson random field analysis which suggests that most polymorphic amino acids in the same proteins are slightly deleterious (1); Clarke B (and also King and Jukes) proposed that at least 85-90% of the mutations causing a change in amino-acid are, on the average, selectively disadvantageous (75,76). For human SNPs, the difference in the number of rare vs. common alleles was used to estimate that 79–85% of amino acid-altering mutations are deleterious (77). The fraction of strongly deleterious mutations (which rarely become fixed) is > 70% in most species, only 10% or fewer of mutations seem to behave as slightly deleterious mutations (SDMs) (65). This maybe the case for RNA virus proteins. For HCV with such a high diversity, there may still be strong purifying selection among most of the sites of its coding proteins. I found that 19% of HCV NS5B aa mutants are accepted mutants which means 81% the mutations are selectively disadvantageous. Hughes AL found that there are Variable intensity of purifying selection on cytotoxic T-lymphocyte epitopes in hepatitis C virus (78). Our results show that there may be strong purifying selection among most of the sites of NS5B protein (and other coded proteins,). Despite the high mutation rate owning to its error-prone RNA polymerase, there is still considerable conservation of virus encoded NS5B protein (and its other encoded proteins, data not shown) among all genotypes found all over the world and there is strong conservative within every genotype and subtype, showing that the selection against deleterious mutations is strong and these purifying selection is the predominant form of selection at the molecular level. Simmonds P proposed that there were constraints on RNA virus evolution at the virus RNA secondary structure level (79). Our study of the nature of factors determining levels its ‘ecological niche’ in the human liver which include the variation of virus proteins also show that there are strong constraints on sequence change of this virus (encoded proteins) both on protein structural and functional and protein stability levels.

## Data Availability

The datasets generated and/or analyzed during the current study are available as Supplementary Data at Zenodo, https://zenodo.org/record/ 5711959, as well as through FTP, https://ftp.ncbi.nih.gov/pub/wolf/_suppl/ virNiches/. Original virus sequences are publicly available for all viruses.

## Acknowledgment

I am indebted to Dr. Minglang He for the suggestion and constructive comments on earlier versions of the manuscript and Qi Dong for assistance in preparing the manuscript.

